# DeepAssembly2: A Web Server for Protein Complex Structure Assembly Based on Domain-Domain Interactions

**DOI:** 10.1101/2024.11.22.624951

**Authors:** Yuhao Xia, Yilin Pu, Suhui Wang, Jianan Zhuang, Dong Liu, Minghua Hou, Guijun Zhang

**Author notes:** Correspondence to Guijun Zhang:* *(G. Zhang).

## Abstract

Proteins often perform biological functions by forming complexes, thereby accurately predicting the structure of protein complexes is crucial to understanding and mastering their functions, as well as facilitating drug discovery. Protein monomeric structure prediction has made a breakthrough in recent years, but the accurate prediction of complex structure remains a challenge. In this work, we present DeepAssembly2, a web server for automatically assembling protein complex structure based on domain-domain interactions. First, the features are constructed according to the input complex sequence and monomeric structures, then these features are used to predict the inter-chain residue distance through a deep learning model, and finally, the complex structure is assembled under the guidance of inter-chain residue distances. Compared with the previously developed version, DeepAssembly2 is trained on a newly constructed inter-chain domain-domain interaction dataset. Meanwhile, several important features have been added, such as Interface Residue Propensity and Ultrafast Shape Recognition. In addition, we introduced the inter-chain residue distance from the AlphaFold-Multimer model to further improve the accuracy. Finally, we also integrate our recently developed model quality assessment method to select the output models. The performance of DeepAssembly2 is significantly improved compared with the previous version, providing a solution for large protein complex structure modeling through divide-and-conquer assembly strategy, and is expected to provide new insights and an effective tool for drug development, vaccine design, etc. The web server of DeepAssembly2 is freely available at http://zhanglab-bioinf.com/DeepAssembly/.

## Introduction

Protein-protein interactions play a vital role in most biological functions. Through interaction, proteins can form complex structures with different morphologies and functions, and intervene in various life processes. Thus, exploring how proteins interact with each other is of great significance for understanding protein functions, revealing disease occurrence mechanisms, and promoting drug design. With the introduction of deep learning technology, protein structure prediction has made great progress in recent years[1-3]. In particular, the emergence of end-to-end protein structure prediction methods represented by AlphaFold 2 (AF2)[4] and RoseTTAFold[5], which have basically solved the problem of protein monomeric structure prediction. While this progress is a major achievement, it is essential to note that accurately predicting the structure of protein complexes remains a serious challenge[6, 7].

For decades, protein complex structure prediction has been a research focus in the field of bioinformatics, and there are numerous methods and tools have been proposed to try to address this challenge. Conventional approaches for predicting the protein complex structure include docking[8, 9] and template-based methods[10-12], which are limited by the accuracy of force-fields and the number of experimentally resolved multimeric structures, respectively. It is impressive that the deep learning-based end-to-end methods greatly improve the accuracy of protein complex structure prediction, e.g. AlphaFold-Multimer (AFM)[13], RoseTTAFold All-Atom[14], and recently AlphaFold 3 (AF3)[15]. These end-to-end models trained specifically on multimeric data from Protein Data Bank (PDB), significantly increase the accuracy of predicted interfaces while maintaining high intra-chain accuracy. Nevertheless, the current end-to-end modeling methods that directly predict structures from amino acid sequences may not be applicable to situations where experimental monomeric structures already exist. This limits its application in practical scenarios such as drug development and vaccine design to a certain extent. Moreover, existing mainstream end-to-end methods may have difficulty meeting the task of modeling large protein complexes due to the limitation of GPU memory. Therefore, it is highly desirable to develop a lightweight method for complex structure modeling through a “divide-and-conquer” assembly strategy[16-18] that takes inter-chain residue interactions as prior information while utilizing known monomeric structures. Furthermore, this method can also effectively utilize the inter-chain interaction information that conform to a specific distribution, which is expected to meet the challenges of multi-conformation or ensemble modeling for protein complexes.

In our previous work DeepAssembly[19] on complex structure modeling, we applied the deep learning network model trained on monomeric multi-domain protein data to the inference of inter-chain interactions of complexes, and regarded the domains in each chain of the complex as assembly units to assemble the protein complex structure. The experimental results in the study show that DeepAssembly can correctly predict the interface, and its prediction results are complementary with AFM. It provides a lightweight way to assemble protein structures by treating domains as assembly units, reducing the requirements for computational resources to some extent. Despite the promising results, the method still has room for improvement in performance, and its overall prediction accuracy still has a gap compared with AFM. The main reason is that although it has been proved that the domain-domain interaction patterns captured from monomeric multi-domain proteins can be used to infer the inter-chain interactions of protein complexes, the domain-domain interactions within the monomer are still limited and cannot fully reflect the domain-domain interaction patterns in a wider range of protein complexes. Therefore, on the basis of the intra-chain domain-domain interactions, it is necessary to further consider the inter-chain domain-domain interactions of the complex, which is the key to improve the performance of the method.

Here, we proposed DeepAssembly2, the latest version of the web server to further improve the performance of complex structure assembly. Compared to the previous version, we constructed a new inter-chain domain-domain interaction dataset to train the deep learning model to learn essential protein-protein interaction patterns. DeepAssembly2 added more crucial features and improved the network architecture to predict the inter-chain residue distance. We also introduced the inter-chain residue distance information from the AFM model and combined it with the predicted inter-chain residue distance to guide the complex structure assembly. Furthermore, DeepAssembly2 integrated our newly model quality assessment method to rank and select the output models. According to the benchmark results, our new version of the DeepAssembly2 web server significantly improves performance over our previous version, and it is competitive with and complementary to other state-of-the-art public servers. DeepAssembly2 will be an important resource for complex structure prediction, and is expected to provide new insights into protein-protein interaction studies.

## Results

### Overview of the protocol

The flowchart of DeepAssembly2 is shown in Figure 1. Starting from the query sequence, the monomeric structure modeling is first performed using the AF2 program. Features are then extracted from sequence, multiple sequence alignments (MSA), and monomeric structures and fed into our in-house inter-chain residue distance predictor, DPIC to generate the inter-chain residue distance map. In parallel, the inter-chain residue distance information is extracted from the AFM prediction model and combined with the predicted inter-chain residue distance to guide the complex structure assembly through a population-based multi-objective optimization algorithm[20]. Finally, the generated models are scored and ranked through our recently developed model quality assessment method, DeepUMQA-X, to output the final complex structure.

**Figure 1.**
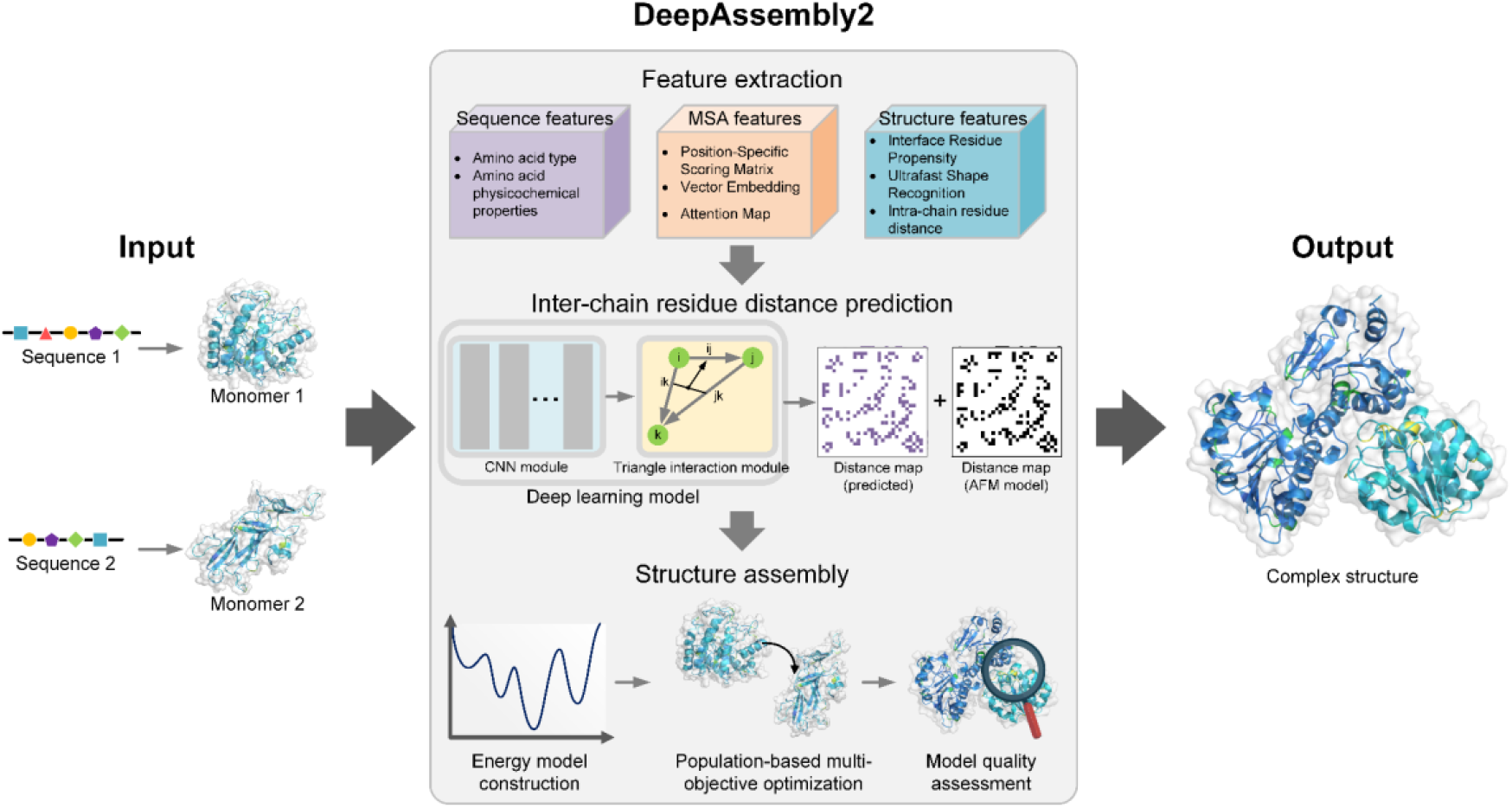
Flowchart of DeepAssembly2. For the input sequence, monomeric structures are first modeled by AF2 program, then features are extracted including three aspects: sequence features, MSA features and structure features. These features are fed into the deep learning model trained on an inter-chain domain-domain interaction dataset to predict the inter-chain residue distance. Next, the complex structure is assembled by a population-based multi-objective optimization algorithm guided by the inter-chain distances we predicted and extracted from the AFM model, respectively. Finally, the output complex structures are selected by using model quality assessment method DeepUMQA-X.

### Benchmark methods

For the benchmark, we compared the prediction performance of DeepAssembly2 with our previous version, DeepAssembly, as well as several other currently state-of-the-art public methods or servers, including AF3, AFM, HDOCK[9], and RoseTTAFold. Specifically, the results of AFM, HDOCK, and RoseTTAFold were obtained by running local stand-alone packages with default parameters, and the AF3 models were predicted by its public webserver (https://golgi.sandbox.google.com/). We used template modeling score (TM-score)[21] and DockQ score[22] to evaluate the accuracy of the built models. The TM-score is a metric defined to evaluate the topological similarity between two complex structures, calculated by the MMalign tool[23]. DockQ measures the interface quality, calculated as:

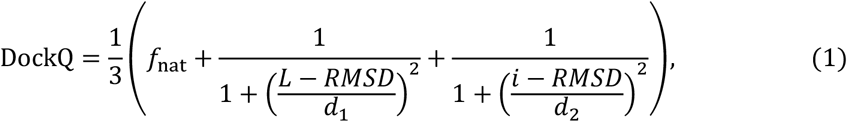

where *f*_nat_ is the fraction of native contacts in the target that is recalled in the model. L-RMSD is the backbone RMSD over the common set of residues of the ligand-protein after the receptor protein has been superimposed. i-RMSD represents the backbone RMSD calculated over the common set of interface residues after these residues have been structurally superimposed. DockQ >0.23 is considered as successfully predicting the interface.

### Performance on the CASP benchmark set

To assess the performance of the server, we collected a benchmark set of 46 complex targets from the 13th to 15th Critical Assessment for Protein Structure Prediction (CASP)[6, 24, 25], including 32 homodimers and 14 heterodimers. The performance of DeepAssembly2 on the benchmark set is shown in Figure 2. DeepAssembly2 achieved an average TM-score of 0.769, a significant improvement over DeepAssembly (0.568) and RoseTTAFold (0.419), and was slightly better than AFM (0.758) and on par with the accuracy of AF3 (0.769) (Supplementary Table S1). Among them, DeepAssembly2 predicted more models with correct global topology (TM-score >0.5) compared with other methods, but the number of models with TM-score >0.9 was less than AF3 (Supplementary Figure S1). In terms of DockQ metric, except for AF3, DeepAssembly2’s average DockQ score (0.439) was higher than that of other methods, and the complex interaction interface was correctly predicted on 56.5% of the targets (Supplementary Figure S2). DeepAssembly2 also performed best in both i-RMSD (7.180 Å) and L-RMSD (16.263 Å), improving by 10.1% and 5.7% over AF3, respectively (Supplementary Table S1). Furthermore, there were a considerable number of DeepAssembly2 models that achieved higher TM-score and DockQ score than AF3 and AFM (Supplementary Figures S3, S4). Overall, the performance of DeepAssembly2 has been greatly improved compared to the previous version, and is comparable to the state-of-the-art complex structure prediction methods, and their advantages will complement each other to a certain extent. Supplementary Figure S5 presents case studies on targets H1143, T1000 and T1179. It can be found that the models predicted by DeepAssembly2 are more consistent with the experimental structures, not only the monomeric structures are accurately modeled, but also the interfaces between the chains are correctly predicted, with DockQ scores of 0.342, 0.573 and 0.339 respectively.

**Figure 2.**
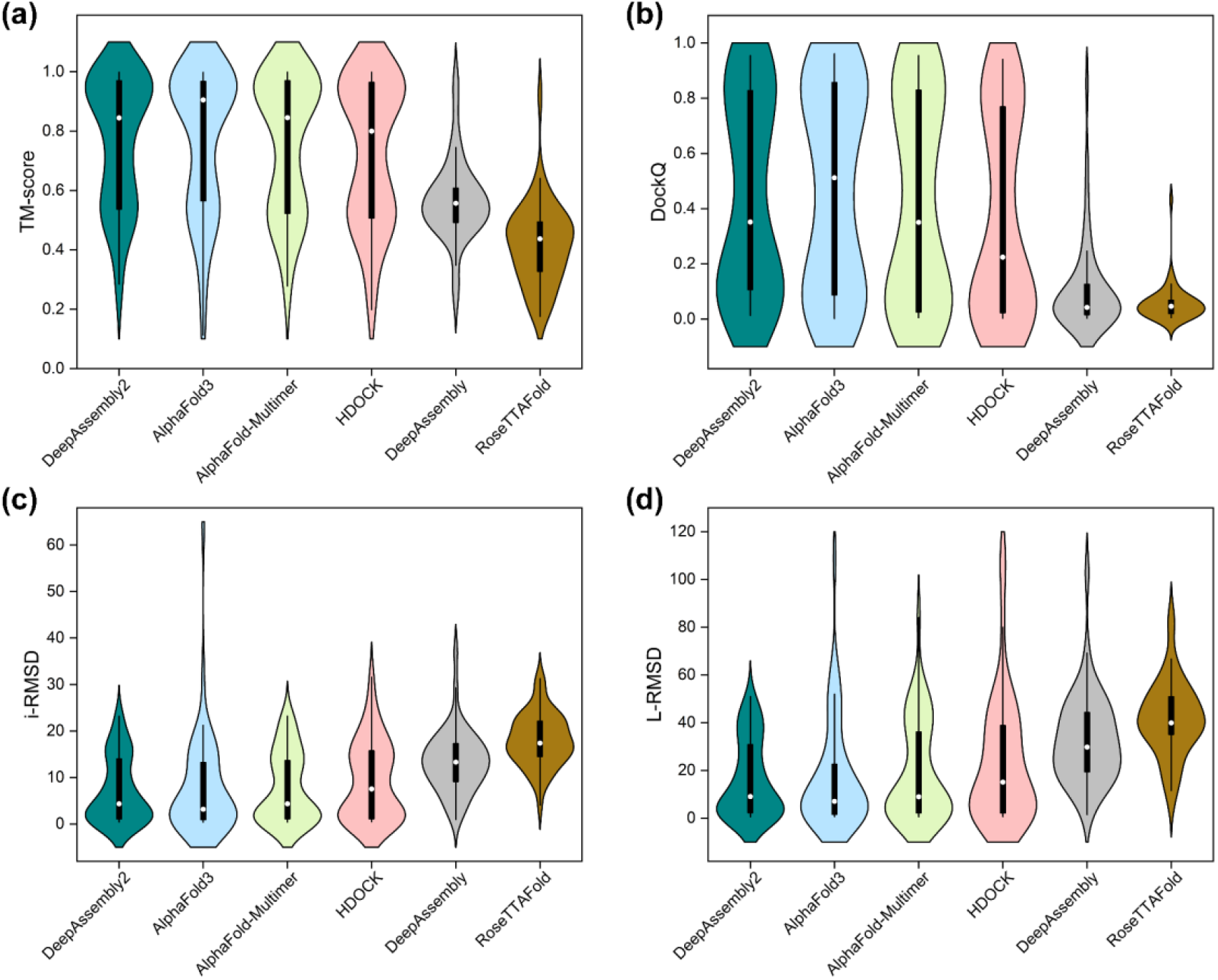
Comparison of the performance of DeepAssembly2 and other methods (AlphaFold3, AlphaFold-Multimer, HDOCK, DeepAssembly and RoseTTAFold) on TM-score **(a)**, DockQ **(b)**, i-RMSD **(c)** and L-RMSD **(d)**.

This improvement is mainly due to the guidance of accurate inter-chain residue distance, which complements the advantages of our predicted inter-chain distance and that from the AFM model. Here, we investigated the contribution of different input features and the dataset used to train the network model to the performance of inter-chain distance prediction. As shown in Supplementary Table S2, the accuracy decreased significantly when the three features were removed respectively, demonstrating the importance of these features in capturing inter-chain interactions. When the constructed inter-chain domain-domain interaction dataset was not used, the performance was also greatly affected, indicating that the domain-domain interaction data can help the network learn the correct inter-chain interaction patterns. In addition to the above, for structure assembly, we also used a multi-objective conformational sampling algorithm to increase the diversity of models and utilized our model quality assessment method, which is also an important factor in improving performance.

### Performance on the heterodimer benchmark set

In general, the structure prediction of heterologous complexes is more challenging than that of homologous complexes due to the relatively less inter-chain co-evolutionary signals. We further evaluated the prediction performance of DeepAssembly2 on the benchmark set of 247 heterodimers constructed in DeepAssembly. Here, we compared DeepAssembly2 with AFM, DeepAssembly and RoseTTAFold, and the prediction results of these comparison methods are from the DeepAssembly paper[19]. On the benchmark set, the average DockQ score of DeepAssembly2 was 0.438, which was higher than other comparison methods, and a 5.5% improvement over AFM (0.415) (Supplementary Table S3). Furthermore, DeepAssembly2 successfully predicted the interface in 166 out of 247 targets, achieving a success rate of 67.2%, and DeepAssembly2 predicted more high-quality models (DockQ ≥0.8) compared to AFM. These results illustrate the advantages of this web server in predicting heterologous complex structures, and it is expected to correctly capture the inter-chain interactions when there is a lack of sufficient co-evolutionary information between chains.

### Webserver interface

The submission and result pages of the DeepAssembly2 web server is shown in Figure 3. The input is the monomeric amino acid sequences (in FASTA format) or monomeric structures (in PDB format) of the protein complex. Users have the option to either input complex sequence/structure data in the designated text box, or upload the FASTA/PDB files. Then, users can submit the task by clicking the “Submit”, and the web server will generate a result page after the task is completed. In addition, if the user enters an email address, they will receive a confirmation email after submitting the task, and an email containing the prediction results after the task is completed. In the webserver result page, it displays the input monomeric sequences and structures, provides the top 5 predicted complex models along with their respective per-residue lDDT and global lDDT scores. Users can download each result individually, or download a compressed package of the full prediction results.

**Figure 3.**
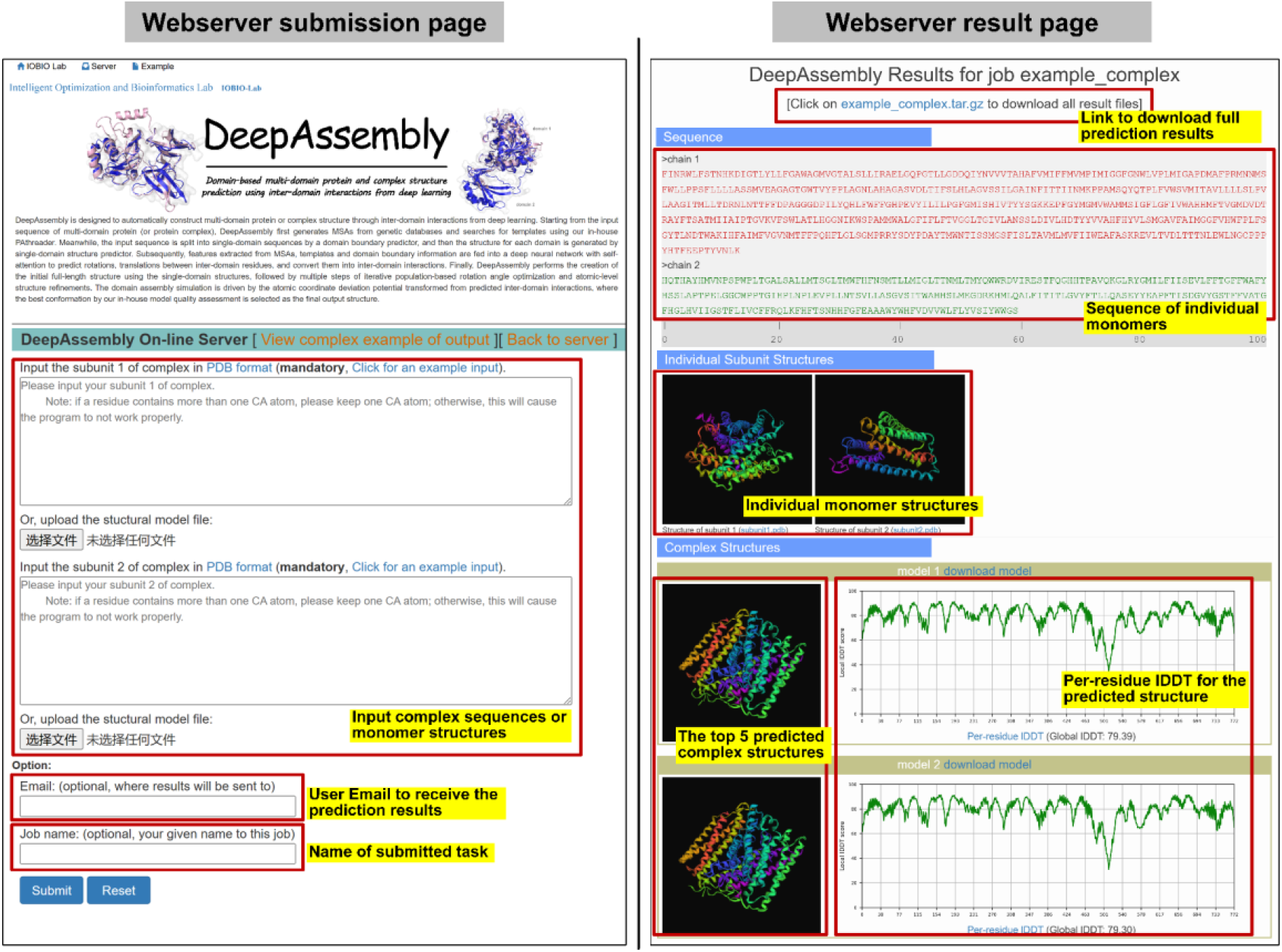
Submission and result pages of DeepAssembly2 web server.

## Discussion

Accurately predicting the protein complex structure is crucial for understanding the protein-protein interaction mechanisms and promoting structure-based drug development. DeepAssembly2 is a significantly updated version of the former pipeline DeepAssembly, in which the performance of protein complex structure modeling has been greatly improved. In DeepAssembly2, the constructed inter-chain domain-domain interaction dataset allows the network to capture essential protein-protein interactions. Additionally, the introduction of new features further improves the accuracy of inter-chain residue distance prediction. The benchmarking performance on complex structure prediction shows that DeepAssembly2 can compare favorably with and complement the current state-of-the-art public servers. It is anticipated that DeepAssembly2 will serve as an indispensable tool for protein complex structure modeling and their interaction mechanism analysis.

## Materials and methods

### Construction of inter-chain domain-domain interaction dataset

We constructed a complex inter-chain domain-domain interaction dataset to train the network model for inter-chain residue distance prediction. Firstly, protein complex structures were distilled from the PDB with the following criteria: (i) entries released before June, 2022; (ii) exclude entries containing DNA/RNA and ensure that only protein structures are included; (iii) the resolution of the structure is better than 3.5 Å; (iv) the biological assemblies contain two or three single chains; (v) the length of the single chain is larger than 20, and a total of 36,955 protein complex structures are obtained. Secondly, all single-chains of these protein complexes were divided into single-domain structures using the DomainParser[26], and the domains belonging to the same protein complex and located in different chains were combined in pairs to form inter-chain domain-domain pairs. Then, we calculated the maximum area of the interface between any pair of domains, and only structure pairs with interface areas >500 Å^2^ are retained. Finally, the remained structure pairs were clustered by MMseqs2[27] with a sequence identity cutoff of 40%, leading to a final set of 16,897 inter-chain domain-domain pairs.

### Description of the input features

The input of the network includes sequence features, MSA features and structure features (Supplementary Table S4).

For the sequence features which contain the amino acid type and physicochemical properties extracted from the sequence. The amino acid type feature is obtained by assigning unique encoding vectors (one-hot encoding) to each of the twenty common amino acids, describing the specificity of amino acids in inter-chain interactions. The amino acid physicochemical properties characterize the polarity, isoelectric point, hydrogen bonding, pH value, etc. of amino acids, which is conducive to the model to identify reasonable inter-chain interaction patterns.

The sequences in MSA reflect the co-evolutionary information of proteins, and its quality directly affects the accuracy of complex structure prediction. For the input complex sequence, we first generate the MSA for each monomer against the Uniclust30_2018_08[28] by using the hhblits[29] (version 3.2.0) with an *E*-value of 1e-3. We then construct the paired MSA according to the organism information[30]. Finally, MSA features are extracted from the paired MSA, including the Position-Specific Scoring Matrix (PSSM), Vector Embedding and Attention Map generated by the pre-trained ESM-MSA-1b[31] model.

Structure features intuitively characterize the spatial structural characteristics of proteins. In order to fully capture the interaction relationship between residues, three protein structure features are extracted from the monomeric structure. The Interface Residue Propensity (IRP) is generated by the PeSTo[32] model, which characterizes the probability that each residue in a monomer forms an interface with other monomers. The Ultrafast Shape Recognition (USR)[33-35] reflects the position of each residue in the overall structure and its adjacent regions, providing information about the relationship between the residue and the overall topological structure. The intra-chain residue distance is defined as the minimum heavy atom distance between pairwise residues within the chain, which provides geometric constraints for predicting the inter-chain residue distance.

### Network architecture

The network architecture for predicting the inter-chain residue distance consists of a convolutional neural network (CNN) module and an attention-based triangle interaction module. For the CNN module which is similar to that used by DeepAssembly, it consists of basic residual blocks. The triangle interaction module consists of triangle update layer, triangle axial self-attention layer and linear transformation layer[36]. The triangle update mechanism updates the pair representations by combining the influence of other residues. It effectively eliminates the inconsistency of pair representations during the prediction process and improves the prediction performance to a certain extent. The triangle axial self-attention layer characterizes the relative strengths of the interaction between pairwise residues through the attention mechanism, and focuses on residues that are highly correlated with the current residue by assigning different attention weights, thereby capturing more accurate interaction information. The linear transformation layer contains two sets of linear transformations, which is used to update the pair representation.

### Inter-chain residue distance energy model

In this work, we used the inter-chain residue distance information to guide the assembly of the complex structure. For the target conformation *x*, we constructed an energy function *E*_*dist*_(*x*) based on the predicted interchain residue distance, which is described as follows:

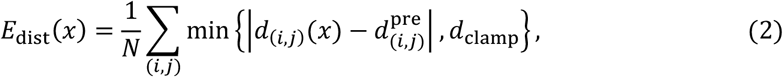

where *N* is the number of inter-chain residue pairs, *d* _(*i,j*)_ (*x*) is the distance between the heavy atoms of the inter-chain residue pairs (*i,j*) for the target conformation *x*, 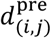 is the predicted inter-chain residue distance, and *d*_*clamp*_ is a clamp, where *d*_*clamp*_ = 30 Å. In order to obtain more diverse prediction models and improve the prediction accuracy, on the basis of our predicted inter-chain residue distance, we additionally introduced the inter-chain residue distance information extracted from the AFM model, and converted it into another energy function 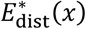, is defined as follows:

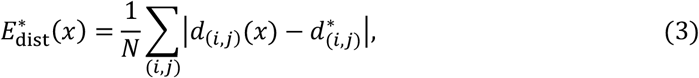

where *N* represents the number of inter-chain residue pairs, *d*_(*i,j*)_ (*x*) is the distance between the Cα atoms of the inter-chain residue pairs (*i,j*) for the target conformation *x*, 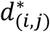 is the distance between the Cα atoms of the inter-chain residue pairs (*i,j*) in the AFM model. In order to combine the advantages of inter-chain residue distance information from different sources, we constructed a multi-objective energy model based on the above energy functions and generated diverse complex structures through a population-based multi-objective optimization algorithm, and the detailed description of the algorithm can be found in our previously published work[20].

In addition, DeepAssembly2 optimized the conformational sampling strategy compared to the previous version. Different from DeepAssembly, which optimized conformation by changing the dihedral angles of residues in the linker region, DeepAssembly2 searched the conformational space by adjusting the rotation and translation between the overall monomeric structures. This allowed DeepAssembly2 to break free from the constraints of the linker during the conformational sampling process, thereby searching a larger conformational space to improve the performance.

### Model quality assessment

For model quality assessment, we recently developed DeepUMQA-X, which used structural consensus to predict quality scores based on global and local perspectives of proteins. From the global perspective, we used different scales to represent proteins and input a multi-level graph neural network to predict the overall quality of the complex. From the local perspective, we used the embedding representation of the protein language model and AF2, supplemented by sequence and structural features, to predict the quality of the local interface score through a graph encoding and decoding module based on an attention mechanism (improved GraphCPLMQA[37]). Further, DeepUMQA-X evaluated all protein models and jointly selected high-quality structures from global and local evaluations as references to predict the quality of complex models through structural alignment.

## CRediT authorship contribution statement

**Yuhao Xia:** Writing – review & editing, Writing – original draft, Visualization, Software, Methodology, Data curation. **Yilin Pu:** – Writing – review & editing, Software, Methodology. **Suhui Wang:** – Writing – review & editing, Software, Methodology. **Jianan Zhuang:** – Writing – review & editing, Software. **Dong Liu:** Writing – review & editing, Software, Methodology. **Minghua Hou:** Writing – review & editing, Software, Methodology. **Guijun Zhang:** Writing – review & editing, Writing – original draft, Visualization, Validation, Supervision, Resources, Project administration, Methodology, Funding acquisition, Conceptualization.

## DECLARATION OF COMPETING INTEREST

The authors declare that they have no known competing financial interests or personal relationships that could have appeared to influence the work reported in this paper.

## Acknowledgements

This work was supported by the National Science and Technology Major Project [2022ZD0115103], the National Nature Science Foundation of China [62173304], and the Key Project of Zhejiang Provincial Natural Science Foundation of China [LZ20F030002].

## Notes

### Competing Interest Statement

The authors have declared no competing interest.

