## Supplementary Material for "DeepAssembly2: A Web Server for Protein Complex Structure Assembly Based on Domain-Domain Interactions"

**Supplementary Table S1.** The performance of DeepAssembly2 and other methods (AlphaFold3, AlphaFold-Multimer, HDOCK, DeepAssembly and RoseTTAFold) for complex structure prediction on the CASP benchmark set. Success Rate (SR) represents the percentage of targets that DockQ score >0.23.

| Method | TM-score | $F_{nat}$ | i-RMSD | L-RMSD | DockQ | SR |
|---|---|---|---|---|---|---|
| DeepAssembly2 | **0.769** | 0.508 | **7.180** | **16.263** | 0.439 | 56.5% |
| AlphaFold 3 | **0.769** | **0.538** | 7.991 | 17.249 | **0.476** | **65.2%** |
| AlphaFold-Multimer | 0.758 | 0.483 | 7.825 | 19.553 | 0.427 | 56.5% |
| HDOCK | 0.731 | 0.399 | 9.304 | 23.516 | 0.374 | 50.0% |
| DeepAssembly | 0.568 | 0.150 | 13.438 | 32.775 | 0.110 | 15.2% |
| RoseTTAFold | 0.419 | 0.091 | 18.076 | 41.647 | 0.058 | 2.2% |

**Supplementary Table S2.** Ablation results of the inter-chain residue distance prediction. "MAE" is the Mean Absolute Error between the predicted distance and the ground-truth distance. "AUC" is obtained by calculating the accuracy of the Top *L* inter-chain residue contact which is converted from the predicted inter-chain residue distance.

| | MAE (Å) | AUC |
|---|---|---|
| Baseline | **5.10** | **0.867** |
| w/o Attention Map | 5.85 | 0.836 |
| w/o IRP | 5.72 | 0.815 |
| w/o USR | 5.49 | 0.816 |
| w/o IRP + USR | 5.99 | 0.854 |
| w/o domain-domain interaction dataset | 5.92 | 0.828 |

**Supplementary Table S3.** The performance of DeepAssembly2, AlphaFold-Multimer, DeepAssembly and RoseTTAFold on the heterodimer benchmark set. Acceptable: $0.23 \leq$ DockQ $< 0.49$, Medium: $0.49 \leq$ DockQ $< 0.80$, High: DockQ $\geq 0.80$.

| Method | DockQ | Acceptable | Medium | High | SR |
|---|---|---|---|---|---|
| DeepAssembly2 | **0.438** | **56** | 67 | **43** | **67.2%** |
| AlphaFold-Multimer | 0.415 | 52 | **68** | 41 | 65.2% |
| DeepAssembly | 0.217 | 21 | 48 | 11 | 32.4% |
| RoseTTAFold | 0.109 | 28 | 17 | 1 | 18.6% |

**Supplementary Table S4.** The input features to the model. $N_{res}$ is the sequence length of the protein complex.

| Type | Feature | Size |
|---|---|---|
| Sequence feature | Amino acid type | [$N_{res}$, 20] |
| | Amino acid physicochemical properties | [$N_{res}$, 7] |
| MSA feature | Position-Specific Scoring Matrix | [$N_{res}$, 20] |
| | Vector Embedding | [$N_{res}$, 768] |
| | Attention Map | [$N_{res}$, $N_{res}$, 144] |
| Structure feature | Interface Residue Propensity | [$N_{res}$, 1] |
| | Ultrafast Shape Recognition | [$N_{res}$, 3] |
| | Intra-chain residue distance | [$N_{res}$, $N_{res}$, 64] |

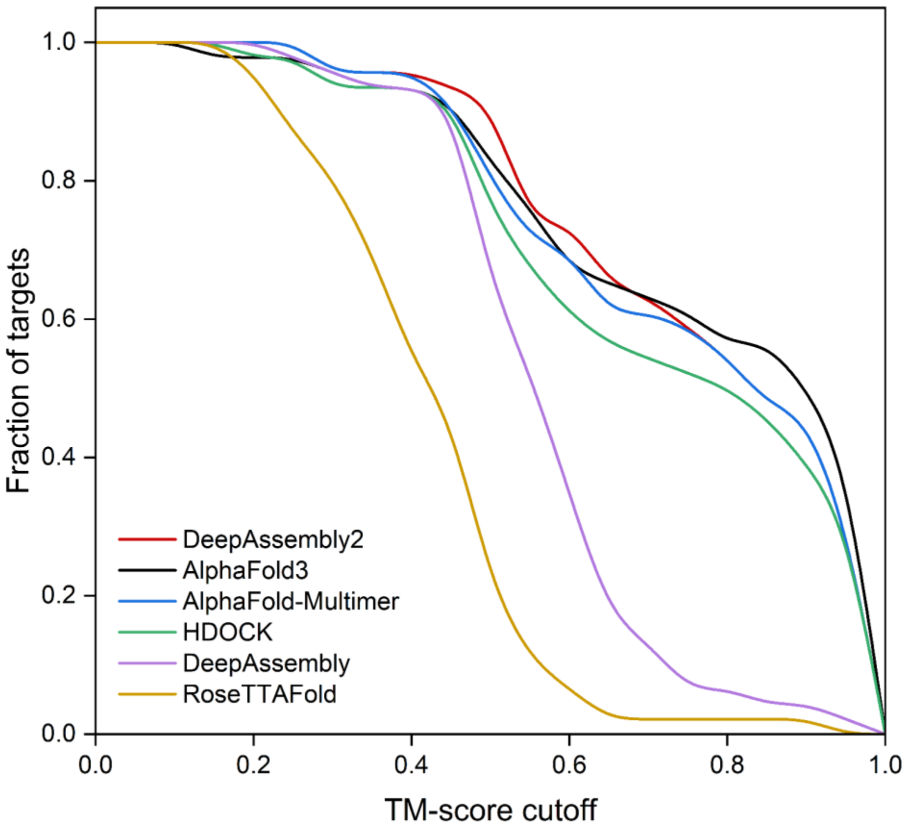


**Supplementary Figure S1.** Comparison between DeepAssembly2, AlphaFold3, AlphaFold-Multimer, HDOCK, DeepAssembly and RoseTTAFold on the CASP benchmark set based on the fraction of targets with TM-score higher than each cutoff.

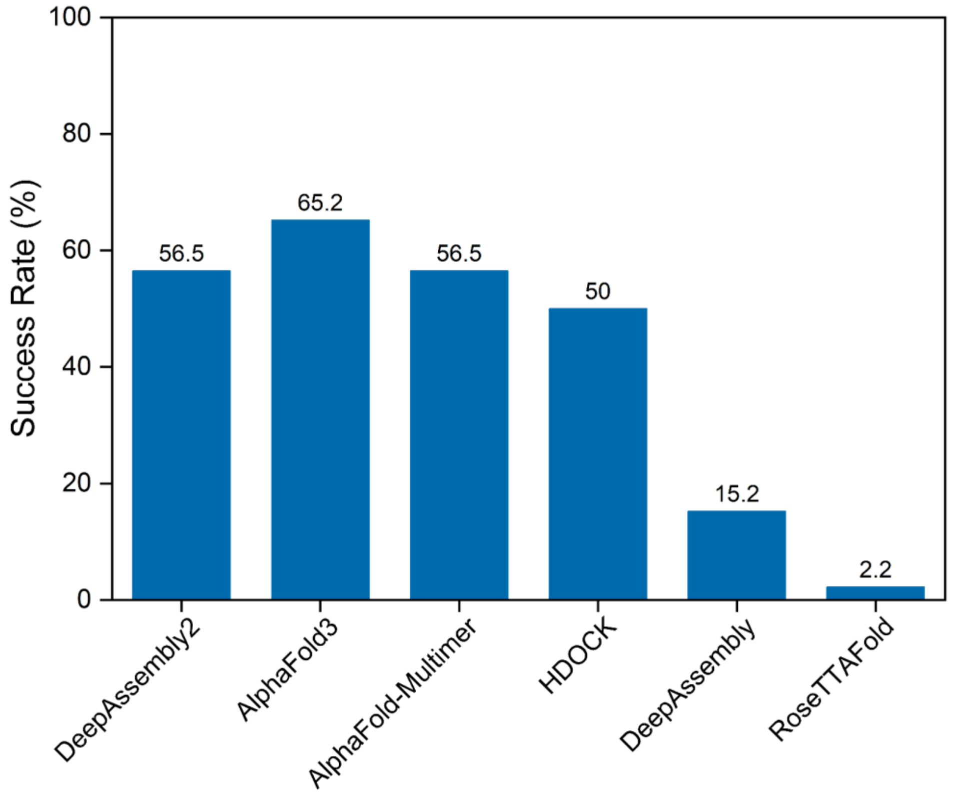


**Supplementary Figure S2.** Success Rate (SR) of DeepAssembly2, AlphaFold3, AlphaFold-Multimer, HDOCK, DeepAssembly and RoseTTAFold in complex structure prediction.

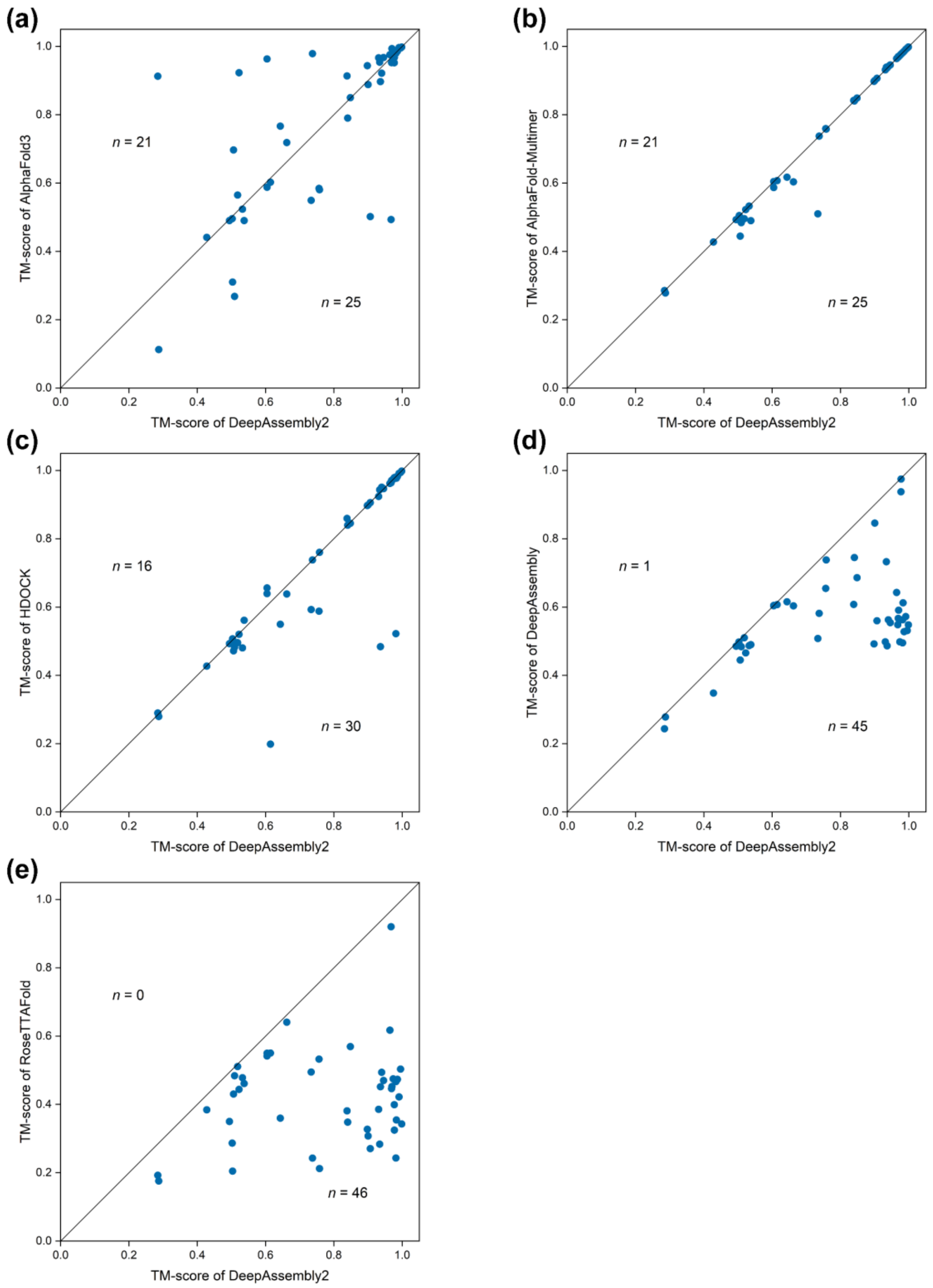


**Supplementary Figure S3.** Head-to-head comparison of the TM-score on each test target between DeepAssembly2 with AlphaFold3 **(a)**, AlphaFold-Multimer **(b)**, HDOCK **(c)**, DeepAssembly **(d)** and RoseTTAFold **(e)**. “n” refers to the number of points on either side of the diagonal.

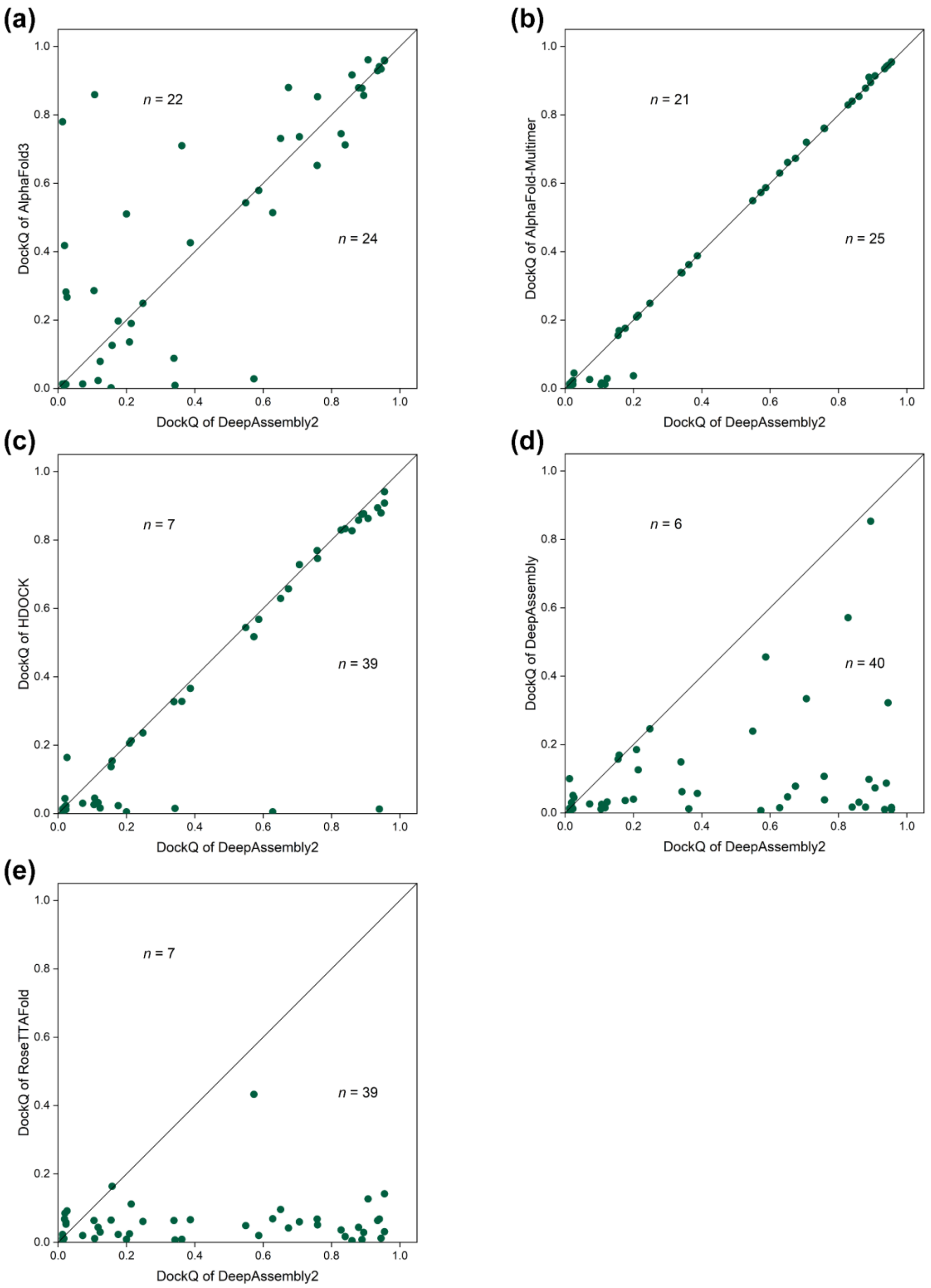


**Supplementary Figure S4.** Head-to-head comparison of the DockQ score on each target between DeepAssembly2 with AlphaFold3 **(a)**, AlphaFold-Multimer **(b)**, HDOCK **(c)**, DeepAssembly **(d)** and RoseTTAFold **(e)**. “n” refers to the number of points on either side of the diagonal.

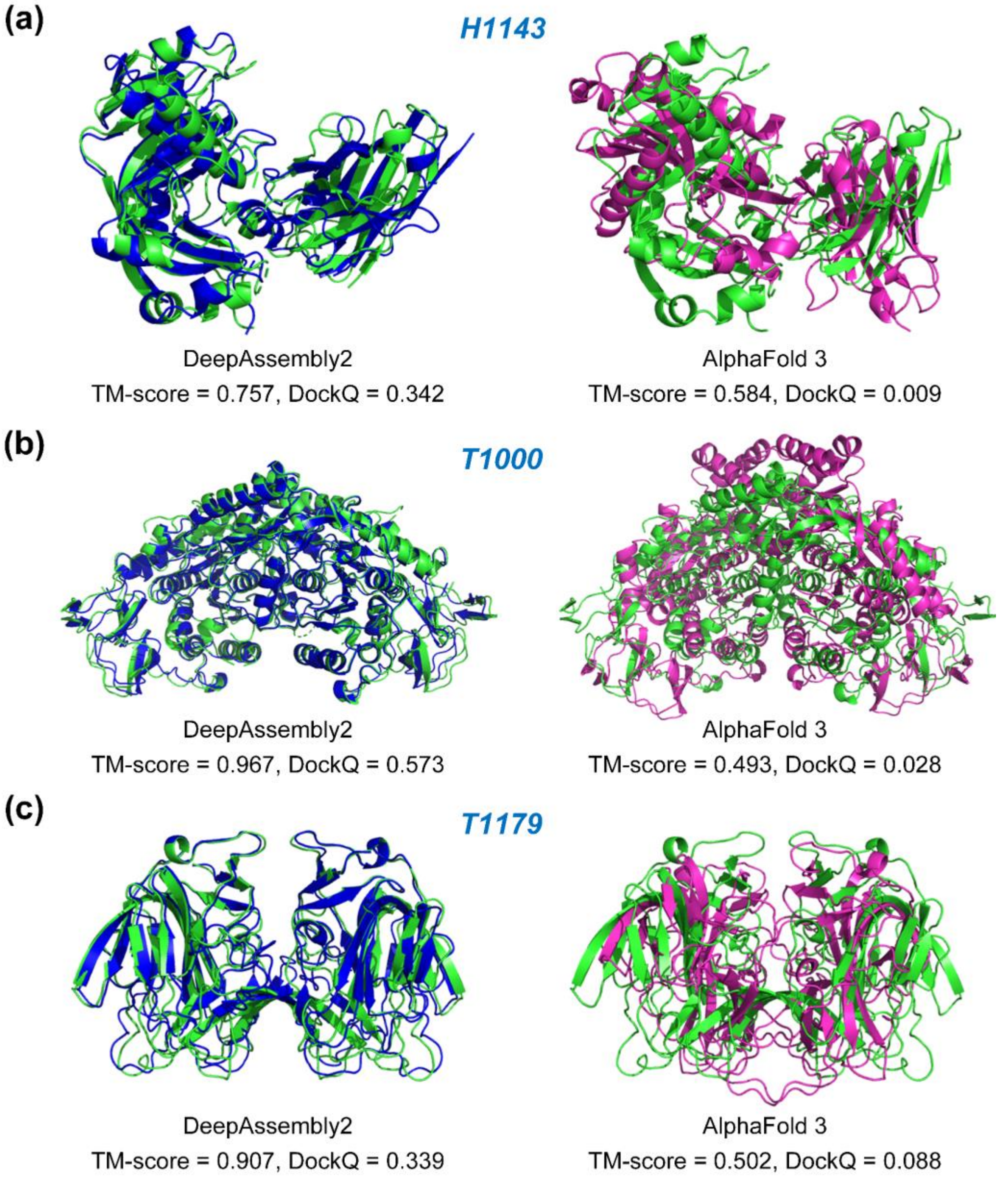


**Supplementary Figure S5.** Case studies of DeepAssembly2 and AlphaFold3 on the CASP targets H1143 **(a)**, T1000 **(b)** and T1179 **(c)**. Experimental structures are colored in green, the models predicted by DeepAssembly2 are colored in blue, and the AlphaFold3 models are colored in pink.
